# A network toxicology approach for mechanistic modelling of nanomaterial hazard and adverse outcomes

**DOI:** 10.1101/2024.01.06.574295

**Authors:** Giusy del Giudice, Angela Serra, Alisa Pavel, Marcella Torres Maia, Laura Aliisa Saarimäki, Michele Fratello, Antonio Federico, Harri Alenius, Bengt Fadeel, Dario Greco

## Abstract

Hazard assessment is the first step in evaluating the potential adverse effects of chemicals. Traditionally, toxicological assessment has focused on the exposure, overlooking the impact of the exposed system on the observed toxicity. However, systems toxicology emphasises how system properties significantly contribute to the observed response. Hence, systems theory states that interactions store more information than individual elements, leading to the adoption of network based models to represent complex systems in many fields of life sciences. Here, we developed a network-based approach to characterise toxicological responses in the context of a biological system, inferring biological system specific networks. We directly linked molecular alterations to the adverse outcome pathway (AOP) framework, establishing connections with toxicologically relevant phenotypic events. We applied this framework on a dataset including 31 engineered nanomaterials with different physicochemical properties in two different *in vitro* and one *in vivo* models and demonstrated how the biological system is the driving force of the observed response. This work highlights the potential of network-based methods to significantly improve our understanding of toxicological mechanisms from a systems biology perspective, guiding the hazard assessment of nanomaterials and other advanced materials.

## Introduction

Chemical safety assessment, comprising the evaluation of exposure hazard and risk, is essential for safeguarding human and environmental health [1]. Hazard assessment relies on model systems to perform an initial evaluation of potential adverse effects of chemicals.

Traditionally, hazard evaluation focused on the compound intrinsic properties, overlooking the context of the exposure that occurs in real-life situations. However, a growing perspective suggests that the molecular configuration of the biological system exposed (here defined as all the active genes and present epigenetic modifications at the physiological stage) can influence the observed phenotype. [2]. This is especially relevant for the emerging class of advanced materials, such as engineered nanomaterials (ENMs), where system-dependent (extrinsic) descriptors take different values depending on external conditions [3]. Indeed, in contemporary toxicology, the observed toxicities are often assumed to be primarily influenced by the dose and the intrinsic properties of the exposure, while the features of the exposed biological system (test system) are referred to as “biological descriptors” of the exposure [4]. However, understanding the system effects is important for selecting relevant models and to correctly characterise the hazard [5]. Thus far, a plethora of omics (e.g., transcriptomics, proteomics) data have been generated to investigate the molecular responses to compounds in various test systems [6]. However, the impact of the biological system on the observed toxicity is frequently overlooked.

The view of living organisms as integrated systems of dynamic and interrelated components has shaped the field of systems biology [7]. This holistic paradigm also includes interactions between organisms with xenobiotics, which is the conceptual foundation of systems toxicology [8]. The systems theory paradigm states that the interactions store more information than the individual elements of the system (*the whole is bigger than the sum^1^*). In this light, molecular profiling of systems alterations are interpreted using analytical approaches with increasing complexity, often based on network theory [9,10]. We previously described a systems toxicology approach for ENM grouping and prioritisation, demonstrating that including molecular alterations of biological systems after the exposure provides better molecular proxies of toxicity [11]. However, traditional approaches in mechanistic toxicogenomics do not explicitly require modelling the interactions between genes, and are based on the assessment of a linear representation of the response [12]. Established methods to characterise chemicals mechanism of action (MOA) using omics data, preprocess and filter data in order to select the most relevant alterations and characterise them functionally. A common strategy uses gene ontology and pathway databases to interpret these alterations [13]. Similarly, other studies interpreted differentially expressed genes and dose responsive genes using the adverse outcome pathway (AOP) framework [4,14]. Such representations, however, are often difficult to associate with system-level effects, and require interpretation and manual reconstruction of the events. We recently demonstrated on a larger set of transcriptomic alterations associated with ENM exposures that, despite the heterogeneity of individual profiles, conserved mechanisms of gene regulation underlie the response to nanoparticulate exposure across multiple species [6]. The patterns of co-regulation can be represented as connections between genes and highlighted by network models. Moreover, we previously used co-expression network inference to characterise the MOA of nanomaterials [15]. However, also in this case, molecular alterations were interpreted *via* functional annotation, requiring the MOA to be manually reconstructed. Therefore, translating gene-level data into interpretable biological responses that could explain the observed toxicity in a mechanistic fashion. Systems effects in toxicology are usually expressed as a causal chain of molecular, cellular and systemic events linking exposures with adverse outcomes (*i.e.*, functional and apical endpoints). These representations are organised in the AOP framework, a multiscale concept connecting early molecular initiating events (MIEs) to adverse outcomes (AOs) through a causally linked chain of events, referred to as key events (KEs) [16]. In hazard assessment, AOPs are usually selected *a priori* based on the end-point of interest. However, the structure of the AOP and the presence of shared events in multiple pathways allows the representation of the entire AOP as a comprehensive network of events which can be applied to a variety of stressors [17–22]. The use of a network of events allows the simultaneous investigation of multiple toxicity mechanisms and the prediction of multiple adverse outcomes. However, limitations still exist in connecting this conceptual framework with molecular alterations (as deduced from omics data). The recently curated molecular annotation of KEs represents an essential step in connecting AOPs to measurable and mechanistic information as derived from toxicogenomics data [14,23].

Here, we describe a novel network toxicology framework that directly interprets the molecular response to chemicals as a chain of toxicologically relevant events. Our framework exploits network properties to contextualise the toxicological responses with respect to the test system. We showcase examples based on a comprehensive set of 31 industrially relevant ENMs of varying chemistries, which are representative of a range of different physicochemical properties and hazard potentials. Our dataset was previously tested on both *in vitro* and *in vivo* test systems, with global expression levels of mRNA, miRNA, and proteins previously assessed from two human cell lines (monocyte-like THP-1 cells and bronchial epithelial BEAS-2B cells), mRNA expression from mouse lung tissues, protein corona profiles, and comprehensive characterisation of all the ENMs [11]. While other studies considering both *in vitro* and *in vivo* assays focused on developing *in vitro*-to-*in vivo* extrapolation models for dose-response evaluation, this study focuses on exploiting network and systems theory to contextualise the molecular response to ENMs with respect to the test system used and interpreting it as a coherent chain of causative events leading to toxicologically relevant effects.

## Results and discussion

### A novel network-based framework links toxicogenomics data to AOPs

Representing the response to an exposure as the result of the biological system-chemical interactions requires modelling approaches that scale from molecular alterations to phenotypic effects in a system-dependent manner. Different biological systems respond differently to the same stimulus, due to their distinct molecular buildup (genotype and epigenotype) [24]. This is reflected by the differences in the expression profiles, both in terms of amplitude of the response, as well as coordinated expression patterns (co-expression) across genes [25]. The response mechanism to an exposure is investigated as all the statistically significant changes at a cellular and molecular level induced by the substance. In transcriptomic studies, this is usually represented as lists of differentially expressed genes (DEGs) which, in turn, are interpreted *via* functional annotation and/or enrichment (Figure 1) [26].

**Figure 1.**
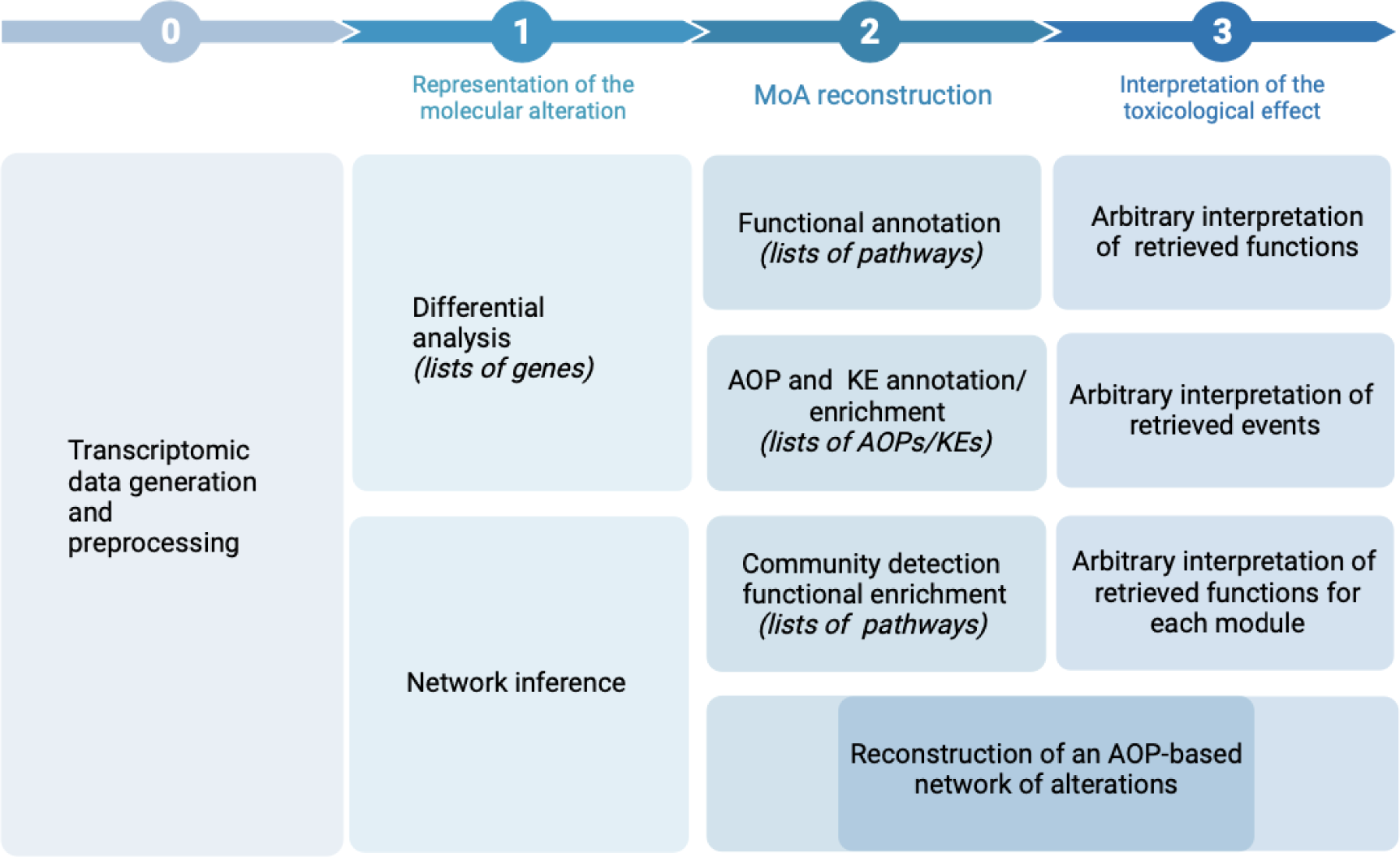
Approaches for the analysis and interpretation of the MOA of ENMs starting from toxicogenomics data. In step 0, toxicogenomics data are generated and preprocessed. In step 1, traditional methods represent molecular alterations in terms of lists of differentially expressed genes (DEGs), while other studies (including this study) model them as co-expression networks. In step 2, previous approaches recontract the mechanism of action (MOA) *via* functional annotation or enrichment of adverse outcome pathway (AOP) associated genes, leading to an arbitrary interpretation of the retrieved functions (step 3). The proposed study allows a reconstruction of an AOP based network of alterations, requiring minimal interpretation with respect to previous approaches.

The obtained results (lists of pathways) require manual interpretation in order to reconstruct the MOA and connect functions with toxicologically relevant endpoints. While previous studies have linked the transcriptomic profile of ENMs exposure to AOP, reconstructing a unique mechanism of response from lists of events still requires manual interpretation. We developed a framework that exploits network properties and the AOP annotation to convert molecular alterations into chains of events across organismal complexity levels (from molecular to tissue/organism). The aim is to interpret input toxicogenomics data exploiting the connection between events (representative of the mechanism) and genes (representative of the test system) in order to reconstruct a model of AOP-based mechanism of action (Figure 2). We tested our framework by characterising the effects of 31 ENMs in three biological systems (the human monocytic cell line THP-1, the human bronchial epithelial cell line BEAS-2B, and lung tissue from C57BL/6 mice exposed by oropharyngeal aspiration). While it is not feasible to characterise the response to all available ENMs in a single study, we selected a dataset as it is representative of different physicochemical characteristics (*e.g,* shape, size, core chemistry, and surface functionalisation) with a range of hazard potentials. [27,28]. This allows to investigate materials with various characteristics, as well as to compare exposures while minimising confounding factors deriving from different experimental designs and nanoparticles manufacturing differences. From the original dataset we identified 93 experimental settings (test system, time, dose, and materials exposed) which we modelled inferring co-expression networks. In this study, an *exposure* will be considered as a combination of these experimental variables. We systematically evaluated the differences arising from the traditional MOA investigation and the use of network-based approaches. All the results for each exposure are reported in the supplementary materials and the comparison results are described in the following paragraph (Supplementary Files, available at https://doi.org/10.5281/zenodo.10390383 in the ‘enrichment_results’ and ‘network_comparison_results’ folders, respectively). In order to identify significantly represented KEs in the exposure co-expression networks, we developed a topological enrichment method. Our strategy identifies KEs as enriched in the co-expression network, if their genes are more densely connected compared to random sets of genes of the same size (cf. Methods) (Figure 2). The enrichment results were then filtered to exclude KEs for which the expression of the annotated genes did not reach a significant alteration, ensuring that the KE had a relevant effect in the biological system under study. The filtered results were then mapped on the AOP network and the MOA was reconstructed. To do so, the most probable MIEs and AOs within the AOP network were selected based on topological and molecular information. The workflow is depicted in Figure 2 (refer to Methods for details).

**Figure 2.**
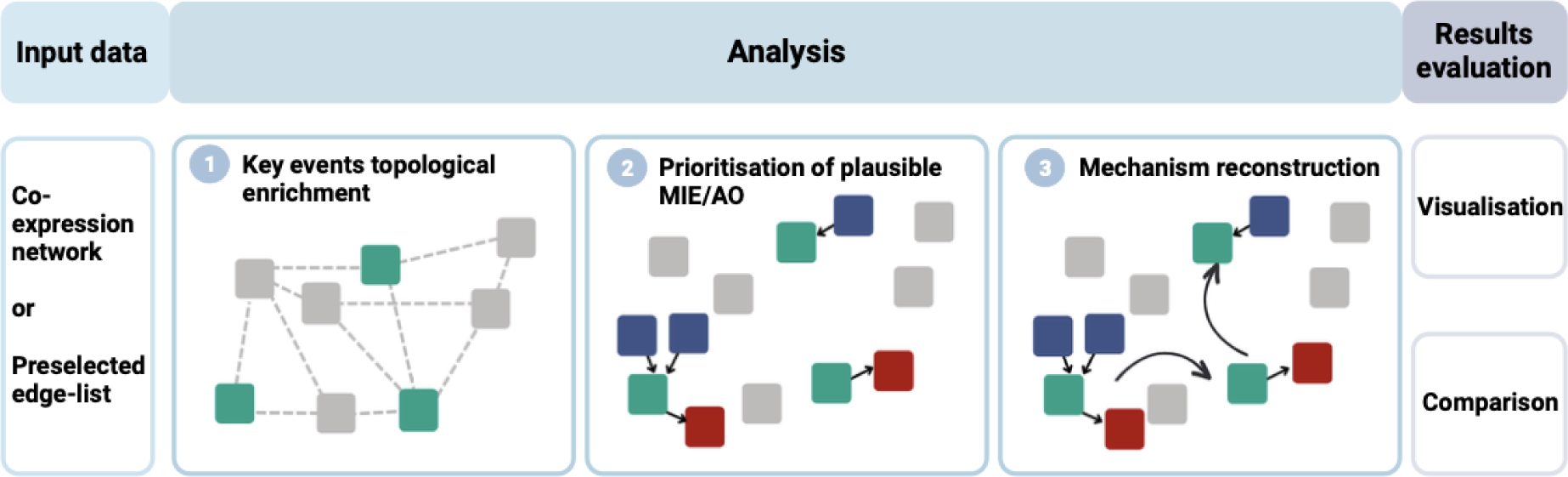
Workflow converting molecular profiles in networks based on the AOP framework. Network models of the exposure (co-expression networks or preselected edgelists) are used as input of the framework. In step 1, a topological enrichment of the adverse outcome pathway (AOP) network is performed based on the input data. In step 2, possible molecular initiating events (MIEs, blue squares) and adverse outcomes (AOs, red squares) connecting to the enriched events (KEs, green squares) are prioritised based on topological properties of the AOP network and molecular alterations. In step 3, the list of MIEs, KEs and AOs is mapped on the AOP network to form an AOP-based interpretation of the mechanism of action (MOA). The results can be visualised as a network or in tabular form, and interpreted independently or compared with other exposure mechanisms.

### Network-based approach can be used to mechanistically interpret toxicogenomics data

In order to analyse the results produced by our framework, we selected three case studies on ENMs exposures which were previously described in [27,28], demonstrating that the proposed approach can recapitulate the main findings while providing additional knowledge with respect to previous findings relying on DEG analysis. Previous *in vivo* studies on the same dataset revealed that the multi-walled carbon nanotubes (MWCNTs) triggered the most prominent changes in the transcriptome amongst all studied materials, and an extensive eosinophil infiltration [28]. Here, we evaluated the reconstructed mechanism of pristine MWCNTs in the mouse lung using as an input the co-expression network derived from exposure data. Our novel network-based framework reported a response mechanism mostly centred on inflammation and reactive oxygen species (ROS) production, which has been largely discussed in the field [29–34]. The enrichment of *“frustrated phagocytosis”* is a relevant event that plays a pivotal role in responses to high aspect ratio materials. For instance, frustrated phagocytosis has been implicated in the prolonged production of proinflammatory cytokines and ROS [35], as well as poor clearance of inhaled particles [36]. All of these factors contribute to the pathogenic potential of MWCNT exposures and favour the development of pulmonary fibrosis, which is a known long term consequence of MWCNT exposure [37]. Our results also highlighted the enrichment of “*decreased fibrinolysis*”, which tightly controls the enzymatic process ensuring the breakdown of fibrin, hence controlling blood clot formation. While the disruption of fibrinolysis is associated with inflammatory processes, direct interactions with ENMs and the coagulation system have also been reported [38,39]. These mechanisms on fibrinolysis may also play a part in the profibrotic effects of MWCNTs [40]. Interestingly, the retrieved mechanism also suggested a substantial epigenetic regulation (ncRNA expression alteration, peroxisome activator receptor promoter demethylation) which has been explored in independent studies on the same material [41,42]. The discussed fibrosis and inflammatory events were well captured by the *in vivo* model, but were less evident in THP-1 cells. Interestingly, only the BEAS-2B cells displayed enrichment for the modulation of the extracellular matrix composition and for alterations of the TGF-β dependent fibrosis pathway. The same dataset revealed the most pronounced inflammation in the CuO-exposed group, associated with high levels of neutrophil infiltration in mouse lungs [28]. Specifically, greater toxicity was observed for the pristine and amino-functionalised forms of CuO [27]. This is also evident in the present study, where the reconstructed mechanisms of response reported apoptosis and decreased cell proliferation in both cell lines. Hence, while the pristine CuO enriched both events associated with inflammation in all the test systems and cell viability, the CuO-NH_2_ showed a massive effect on cell proliferation in the two cell lines, and impact on the DNA repair, chromosome stability and altered microtubule dynamic in the *in vivo* model. Interestingly, the predicted MIEs included alteration of the oxidative status in all the test systems, and alteration of ion pumps and receptors in both lung and BEAS-2B. This is in line with the other findings that report a toxicity mechanism mainly based on oxidative insults [43]. Similar results were also observed in the A549 cell line, revealing an acute genotoxicity of nano-sized CuO, which is well captured by our framework [43–45].

We also investigated the response mechanism to titanium dioxide (TiO_2_) in the mouse lung, and compared the effects of variations in the material geometry (Figure 3). Previous studies demonstrated that the spherical TiO_2_ or rod-shaped induce different responses, albeit without describing the specific mechanism [28]. Our framework highlights the effect of the rod shape on the cell membrane, with initiating events comprising narcosis and decompartmentalisation (Fig 2B). Alterations of the cell membrane also affect lipid composition and induce the activation of AKT2 signalling, which regulate many processes including metabolism, proliferation, cell survival, growth and angiogenesis [46]. The spherical TiO_2_, instead, showed its main effect on inflammation, ROS production, ion balance alteration, and effects on the calcitonin gene-related peptide (CGRP), inducing respiratory alterations (Fig 2A and Supplementary Files, available at https://doi.org/10.5281/zenodo.10390383 in the ‘enrichment_results’ folder). Moreover, the framework captured the genotoxicity associated with TiO_2_ nanoparticles, one of the concerns behind the recent and much-debated EU ban on their use as food additives [47]. The effects of rod-shaped TiO_2_ have been scarcely described to date; however, other studies on rod-shaped nanocarriers and rod-shaped silica and silver nanoparticles reported a significant effect of shape, leading to increased cell uptake, especially by epithelial cells [48–50]. Notably, BEAS-2B response mechanisms to TiO_2_ showed substantially more alterations with respect to the other test systems. In sum, we have showcased a selection of the results, using MWCNTs, CuO, and TiO_2_ as exemplars, to demonstrate the potential of our framework and how it can alleviate the problem of toxicogenomics data interpretability when profiling the molecular response to chemical exposure.

**Figure 3.**
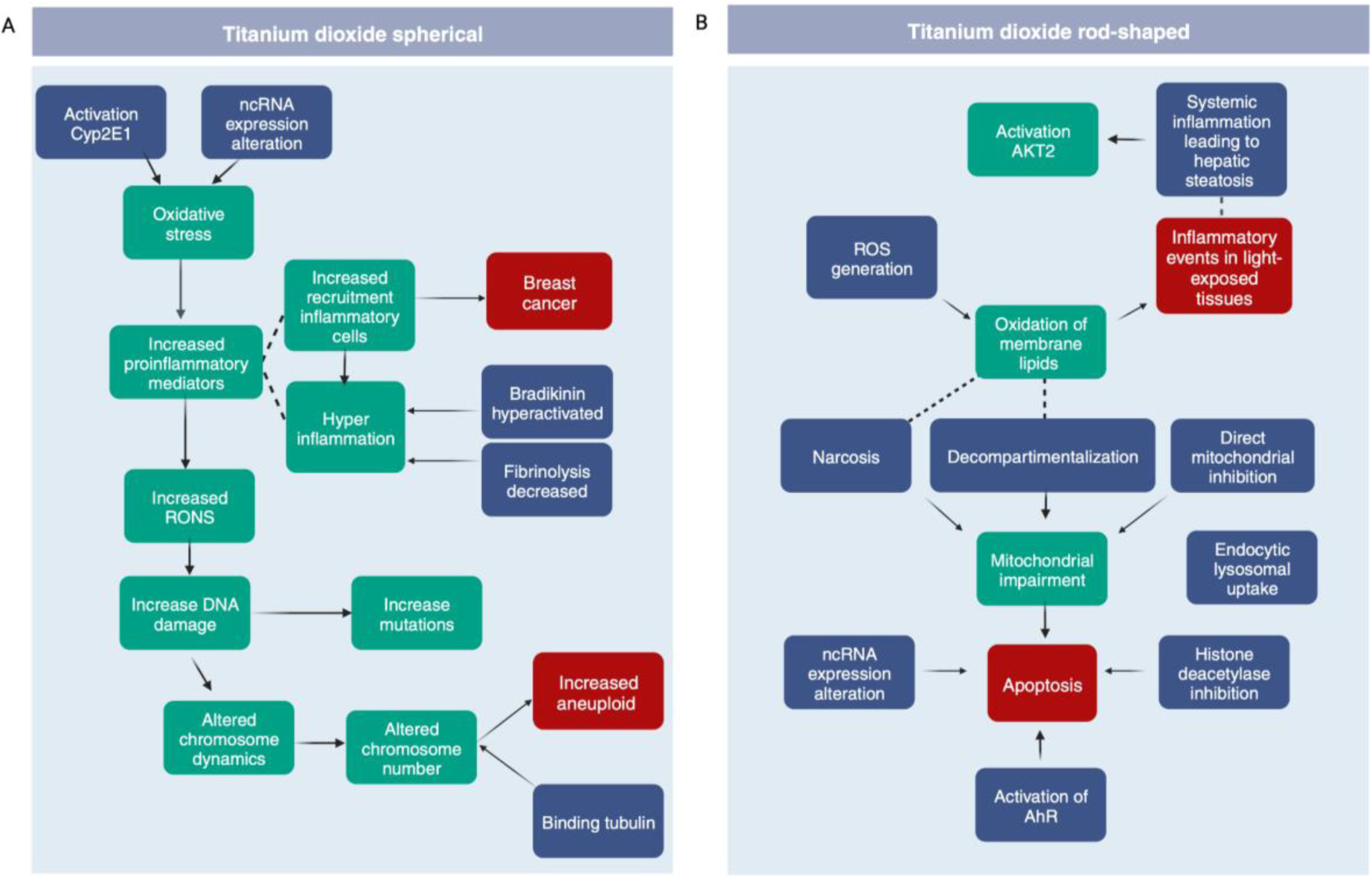
Reconstructed mechanism of response to titanium dioxide (TiO_2_) exposures with varying shapes, i.e., spherical (A) versus rod-shaped (B). Enriched key events (KEs), molecular initiating events (MIEs) and adverse outcomes (AOs) are reported in green, blue, and red, respectively. Dashed lines connect events which are not formally linked in the AOP-Wiki database, but are related to common biological processes (not provided as output of the framework).

### Representing the MOA as a network increases the information content

Systems biology relies on the principle that connections store more information than isolated pathways or gene sets [12]. We previously demonstrated that different transcriptomic profiles in response to nanoparticulate share similar regulatory mechanisms [6]. Gene-to-gene connections can be modelled through the construction of co-expression networks, where genes are linked based on the presence of a statistically significant co-expression. These relationships may originate from shared functional attributes, involvement in common pathways, participation in protein complexes, or engagement in transcriptional regulation [51]. Therefore, we tested whether the response mechanism extracted from co-expression networks *via* our framework would provide better qualitative and quantitative results than the one relying solely on gene sets.

In order to evaluate how effectively the network based strategy performs with respect to the traditional representation of ENM responses (Figure 2), we compared the detected mechanisms when levels of neutrophil infiltration induced by ENM exposure are compared [11]. To ensure an unbiased comparison, we used as input our framework lists of DEGs and performed a classic enrichment of the KE, and compared the retrieved mechanisms. In both cases, our framework detects inflammatory events associated with exposures with low neutrophil infiltration. Early pathological alterations in the lung are usually linked to injury of pulmonary endothelial cells and vessel destabilisation [52]. The network-based approach did capture this element, together with TGF-β activation, when medium neutrophil infiltration is induced. The gene set based approach, instead, captures the TGF-β activation for high infiltration levels only. Interestingly, exposures resulting in high neutrophil infiltration induce “*oxidation of membrane lipids*” and “*NRF2 depression*” in network reconstructed mechanisms, but not in the ones derived by gene sets. NF-E2-related factor-2 (Nrf2) is a regulator of cellular antioxidant responses, largely expressed by neutrophils and only recently identified as a key player in the regulation of inflammasome activation [53]. The coexistence of oxidative stress markers, lipid peroxidation, inhibition of NFkB, and a high neutrophil infiltration (as evidenced by the network approach) is well known as being prognostic of chronic and deregulated inflammation and has recently been associated with lung function deterioration in COVID-19 [54].

In order to produce a quantitative evaluation of the use of network models to represent exposure responses, we defined measures of information content and compared the results obtained from gene sets and co-expression networks based strategies (Figure 4). Detailed results of the framework for each set of DEGs is reported in the supplementary files (available at https://doi.org/10.5281/zenodo.10390383 in the ‘network_comparison_results’ folder). When profiling the MOA of an exposure, information gain can be defined as the ability to reconstruct a complete mechanism connecting the initiating event(s) to observable outcome(s). In this light, we defined the “AOP completeness score” as a measure of the ability to observe a complete chain of events connecting one or more MIE and one or more AO (Figure 4A, and refer to Methods). A desirable gain can also be expressed in terms of less uncertainty of the observed mechanism, indicating a more predictable or homogeneous exposure response. Therefore, we defined the “Key event connectivity score” as the ratio between the portion of connected KEs against the unconnected ones characterising the MOA of an exposure. For the same exposure, the DEGs and the co-expression networks were used as input to obtain two independent models of the MOA. In both cases, we demonstrated that both the “AOP completeness score” and the “Key event connectivity score” showed higher values when a co-expression network was used as input (Figure 4, Supplementary Files available at https://doi.org/10.5281/zenodo.10390383 in the ‘network_comparison_results’ folder). Taken together, these results demonstrate that a network-based representation of toxicogenomics data systematically outperforms a traditional analysis based on the evaluation of individual molecules. For this reason we focus on the results obtained from co-expression networks only.

**Figure 4.**
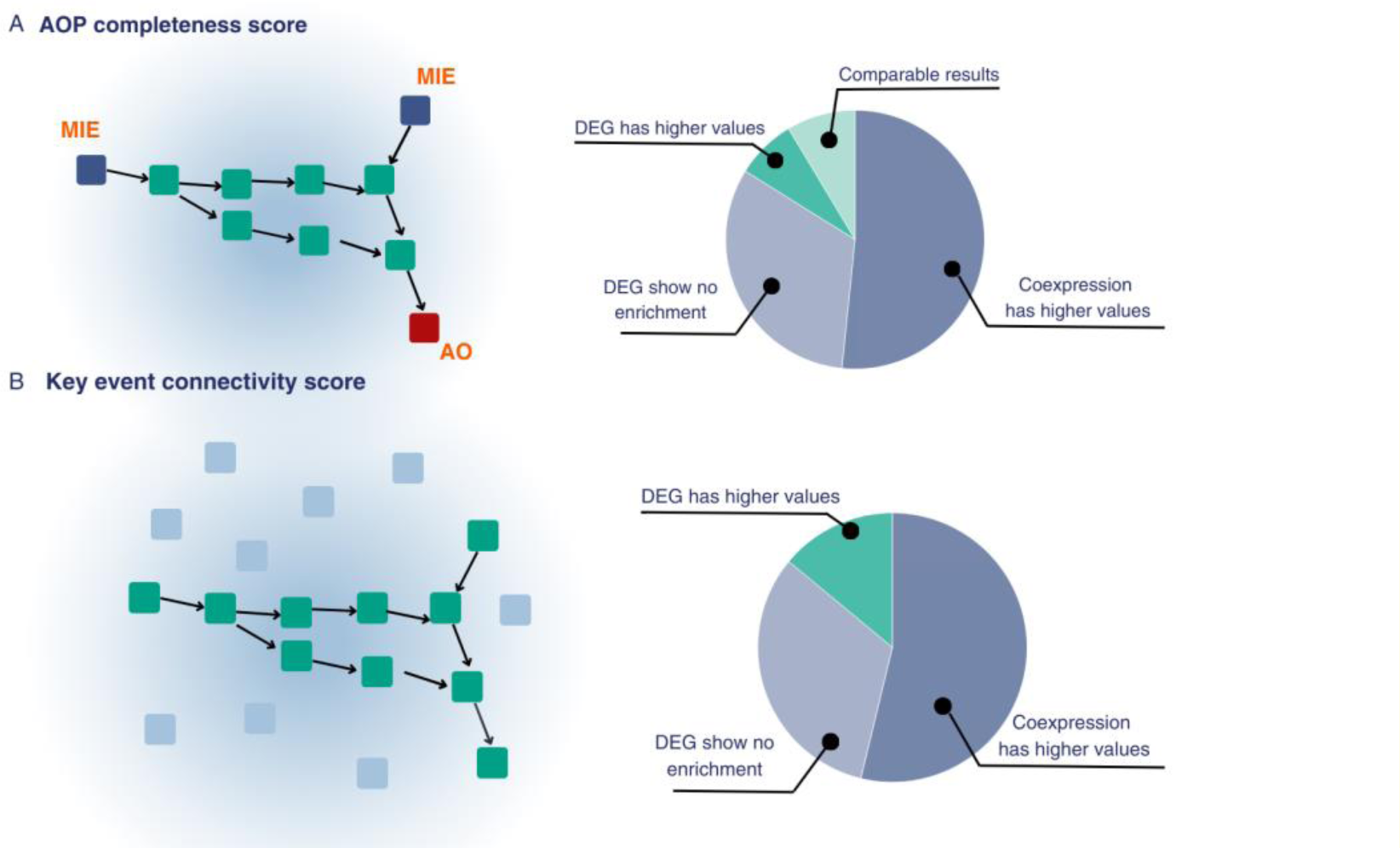
Graphical representation of the defined information score based on the possibility of reconstructing the mechanism of action (MOA) of the compound. A) The “AOP completeness score” was defined as a measure of the ability to observe a complete chain of events connecting one or more molecular initiating events (MIEs) and one or more adverse outcomes (AOs). B) The “Key event connectivity score” was defined as the ratio between the portion of connected key events (KEs) against the unconnected ones characterising the MOA of an exposure. For each score, pie charts representing the results in the information metric values when mechanisms of response are reconstructed based on lists of differentially expressed genes (DEGs) and co-expression networks.

### Network-based models of exposure highlight mechanisms related to different hazard levels

A current need in toxicological assessment is to prioritise highly hazardous materials for comprehensive testing. However, previous efforts showed that toxic responses cannot be clearly distinguished when analysing whole transcriptomic profiles or by relying only on specific physicochemical properties [55]. Given the increase in information content of the network approaches, we investigated whether it was possible to identify differences in nanomaterials exposures with varying toxicity levels. First, we grouped the 93 exposures based on the toxicity endpoints we had collected in [11], which are based on an integration of cyto-, geno-, and immunotoxicity data. Second, for each group, we extracted overrepresented edges in the co-expression networks of the same group. This allows the extraction of features which are representative of that group, but not of others. Finally, the edges were interpreted in the AOP context using our framework. Our results showed that low hazard exposures (as defined in [11]) were associated with oxidative stress, recruitment of inflammatory cells, and decreased lung function. In intermediate hazard exposures, cell proliferation alterations emerged, as well as cell activation, and cytoskeleton modifications. Among the most relevant MIEs represented in medium hazard nanomaterial exposures, we found frustrated phagocytosis and tubulin binding. Previous studies on high aspect ratio materials proved that mesothelial cells initiate pro-inflammatory responses when retaining materials, even after weeks, and can play an important role in stimulation of tumour growth [56,57]. Highly hazardous materials mainly enriched adenomas and carcinomas as AOs. Interestingly, the relative representation of AOs was the highest in the events enriched by highly toxic compounds, with the amount of molecular initiating events following the opposite trend (Figure 5). This suggests that a possible difference between low, medium and highly toxic materials is related to the induction of alterations which are indicative of an adverse outcome. It is known that different stimuli can alter a cell state either in an irreversible or reversible manner [24]. In the first case, the molecular response facilitates the transition of the cell from the steady state state to a new stable state, whereas in the second case a temporary unstable state is reached that may be reverted to the initial one, once the stimulus is removed [24]. In the context of an AOP, reaching an adverse outcome represents a multiscale event implying that alterations have happened across biological levels of organisation (molecular, cellular, tissue, organism). Tissue responses, for example, underlie cellular and molecular complex events that need to happen in order for the macroscopic effect to emerge. Our hypothesis echoes the “hierarchical oxidative stress model” of Nel *et al.*, where the nanoparticle capability of inducing different levels of ROS, triggers molecular and cellular events with increasing toxicity potential [58]. Our results suggest that highly hazardous materials may induce an alteration that quickly spreads across biological levels and moves towards a new phenotypic state which is, therefore, harder to revert.

**Figure 5.**
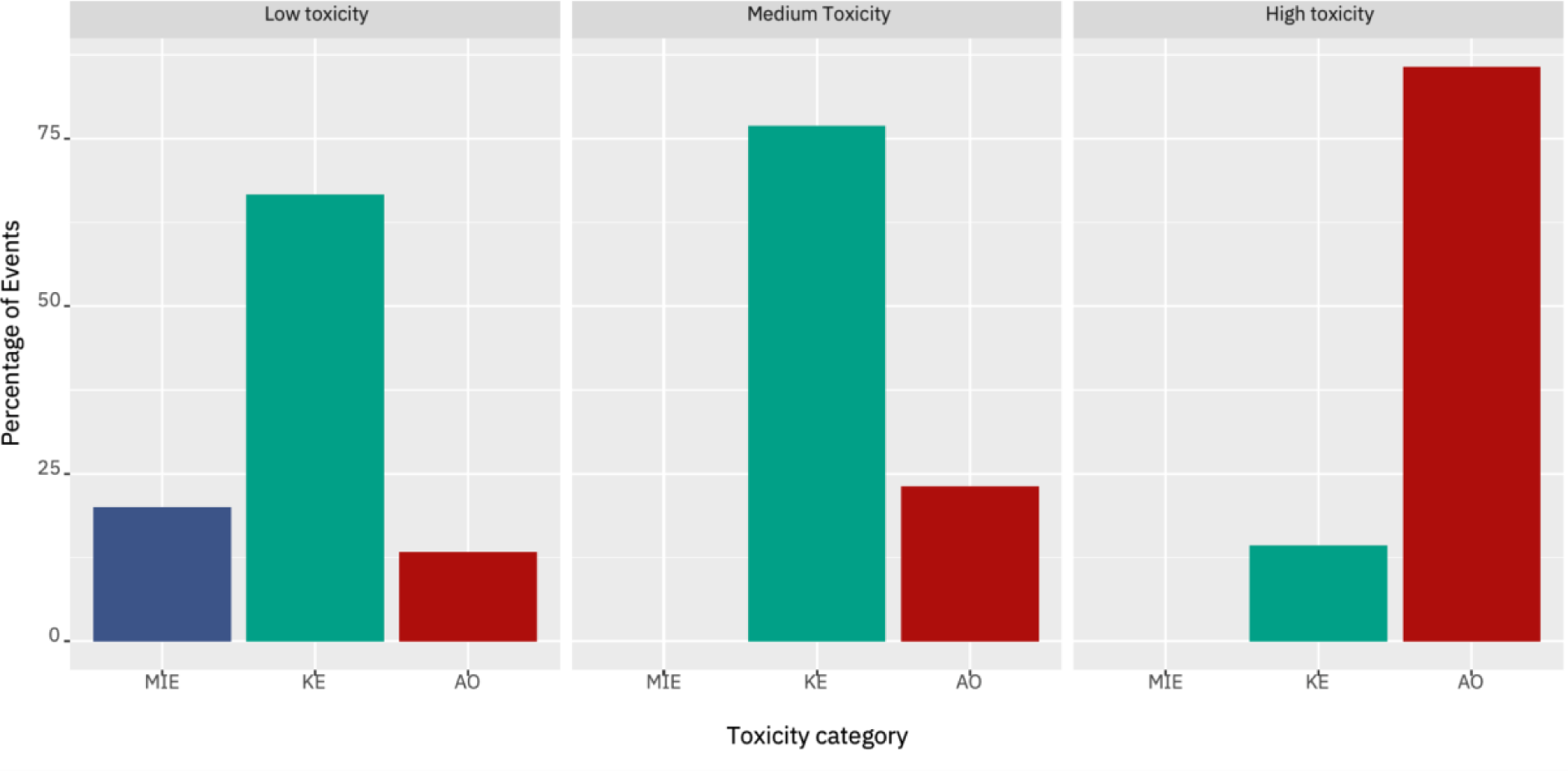
Distribution of molecular initiating events (MIEs), key events (KEs) and adverse outcomes (AOs) across engineered nanomaterials (ENMs) toxicity classes (low, medium, high). Data are reported as a percentage of the total enriched events.

### Network properties underline the impact of the exposed biological system

The results discussed above pointed towards differences between the test systems. Therefore, we decided to investigate the overall impact of the test system on all the 93 exposures we previously defined (i.e., dose, time, materials, and test system) by clustering the inferred co-expression networks based on their topological properties. Our analysis showed that biological systems have a significant impact on the response to ENM exposure, with exposures to different materials in the same test system clustering together (Figure 6A). Furthermore, exposures in the same biological system share portions of the response identified as the set of statistically overrepresented edges in the networks inferred from the transcriptomics data profiled in the same biological system. These overrepresented edges underlie functions which are representative of the system under evaluation (Figure 6C-E), such as inflammogenic functions for THP-1 cells (Figure 6C), phagocytosis and endocytosis for BEAS-2B cells (Figure 6E). However, for mouse lungs, this behaviour is less evident, with less overrepresented edges and functions shared across the exposures. This is possibly due to the cytological complexity of the tissue [59], as well as to the different growth conditions of the test system which may affect the biocorona formation on ENMs [59] (Figure 6D).

**Figure 6.**
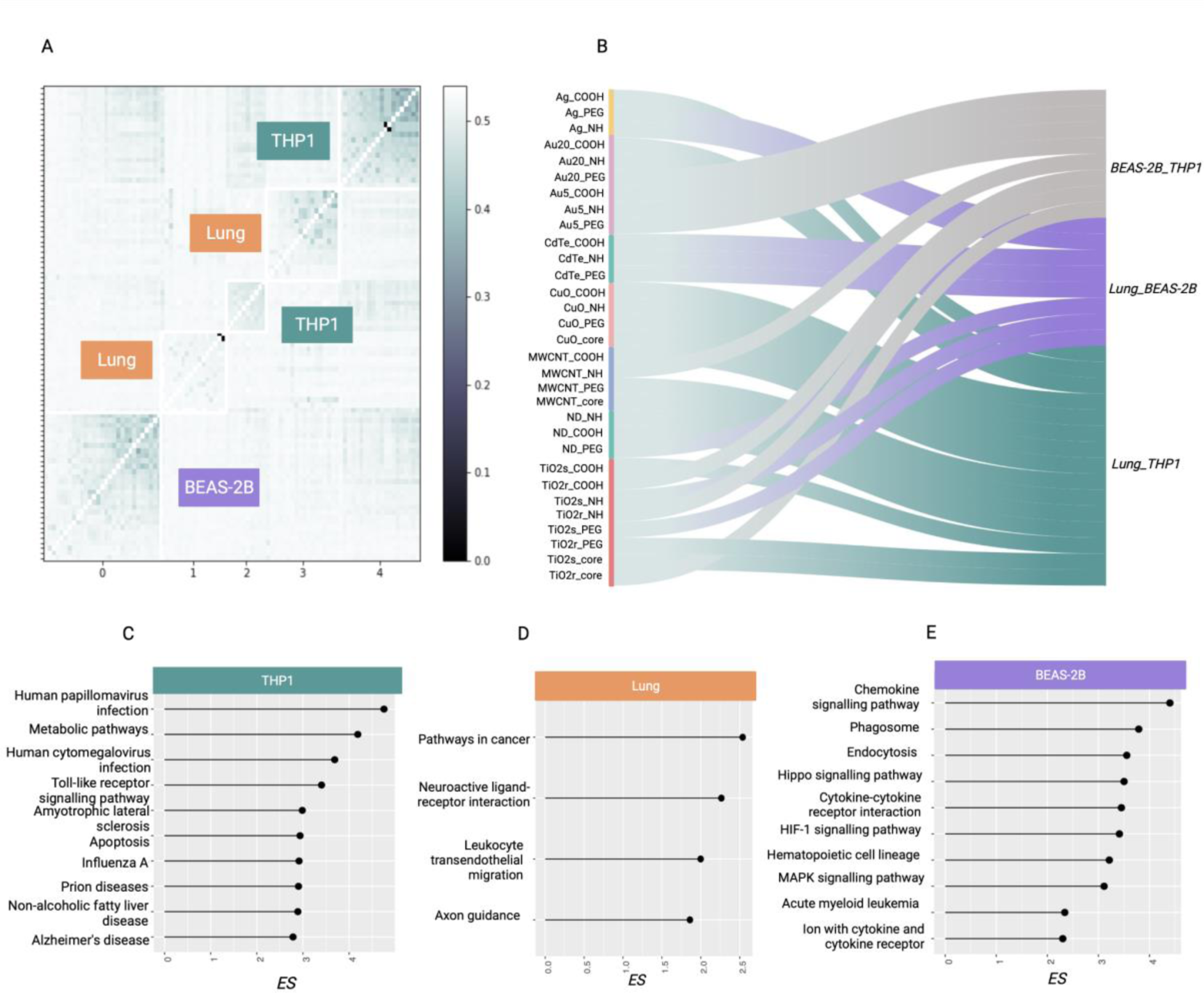
Biological systems influence the response to ENM exposures. A) Clustering of the 93 exposures co-expression networks based on topological properties (edge betweenness, Jaccard, Hamming distance, SMC distance between the existing edges of the networks as well as a percentage value of shared edges between each pair of input networks). B) Similarity between biological systems in exposures of the same material. In teal, exposures where the highest similarity is between lung and THP-1, in violet when the highest value is between BEAS-2B and lung, in grey when the *in vitro* systems are not representative of the *in vivo* counterpart. Engineered nanomaterial (ENM) chemistry has been annotated. C-E) Top fifteen enriched pathways of the overrepresented edges for each biological system, with their respective enrichment score (ES).

In order to evaluate the impact of the biological system on the molecular response, we clustered the reconstructed AOP-based response mechanisms for each exposure. We observed that the exposures cluster based on the induced biological response, rather than the exposed material. Our results indicated that even when exposed to the same ENM, the observed response mechanism is a function of the interaction between the compound and the test system. Indeed, the 93 exposures clustered in two groups, almost completely separating the THP-1 and BEAS-2B exposures. When a major inflammatory response was generated, THP-1 responses were closer to the responses in mouse lungs. On the contrary, when cell stress responses, mainly based on DNA damage, were activated, exposures in BEAS-2B overlapped more with the lung mechanism. Lung exposures split in the two groups according to the nature of the tissue response. These results do not imply the validity of the in vitro or in vivo models. Indeed different models provide different insights in the MOA of chemicals, which need to be considered when evaluating the results.

Several previous efforts have focused on the possibility of identifying physicochemical similarities between the ENMs to predict common responses [60]. Our results suggest that *in vitro* test systems, due to their molecular buildup, usually show only portions of the complete response, as already described [14]. On the other hand, in complex systems, such as the lung tissue, a core mechanism drives the response, but it is difficult to dissect the contribution of specific cytological components. This supports the idea that, in order to correctly estimate hazards, information on the test system must be considered, as the observed mechanism of action depends on the test system used for the assessment even in comparable experimental settings (same material, same or equivalent doses and time of exposure). To further investigate this phenomenon, for each ENM in the original dataset, we compared the similarity of the *in vitro* systems and the *in vivo* counterpart. This is an important step in light of efforts to develop alternative methods to animal testing, where the *in vivo* system is often considered as the benchmark for new approach methodologies (NAMs). Other investigators have compared animal experiments with non-animal-based alternatives in order to assess *in vitro*-to-*in vivo* extrapolation (IVIVE) [61]. However, defining IVIVE relationships requires a focus on doses and physiologically based kinetic models, which is out of the scope of this study. Here, we focused on the impact of the *in vitro* or *in vivo* test system with respect to the observed response. In order to exploit the network properties to assess the similarities or differences between *in vitro* and *in vivo* systems, we defined a “biological system similarity score” as the comparison between the response mechanism in pairs of systems, with respect to the complete AOP network (refer to Methods for further details). More than half of the ENMs showed a higher similarity between mouse lungs and THP-1 cells, while the remaining exposures equally split in the other two categories representing mouse lungs and BEAS-2B cells, and THP-1 cells and BEAS-2B cells, respectively (Figure 6B). This is not unexpected, as many ENMs induce an inflammatory response that is well captured by cell lines such as THP-1 [62], suggesting that specific properties of the cell lines allow the mechanistic translatability across test systems. However, in 25% of the cases, the THP-1 and BEAS-2B *in vitro* systems shared more similarities than the *in vivo* system, suggesting that the *in vitro* assessed hazard was less overlapping with tissue-level observations in one quarter of the exposures. Thus, selecting the appropriate cell line(s), or utilising a panel of cell lines of varying origin, may be important [2,63].

### The molecular buildup influences the applicability domain of hazard tests

Given the remarkable impact of the test systems on the observed response mechanism, we investigated whether the molecular buildup of the biological system would affect the assessable hazard with respect to the complete set of KEs in the AOP framework. An AOP represents the toxicologically relevant events to be tested in order to mechanistically assess the hazard of chemicals. To this aim, we combined the AOP-based response mechanisms of all 93 exposures in the different test systems, and represented them as a portion of the AOP network (Figure 7).

**Figure 7.**
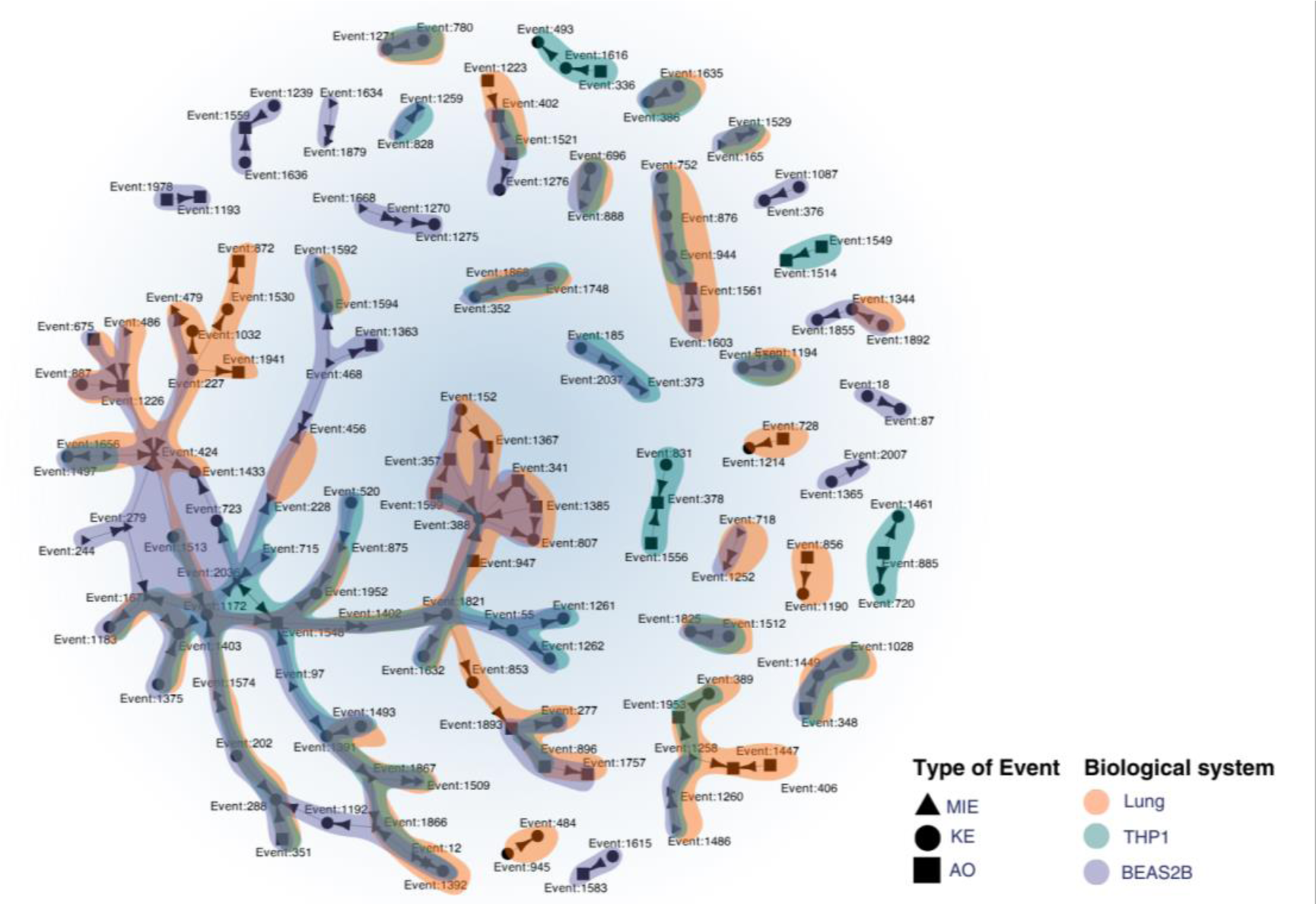
Explorable AOP network is determined by the biological system. Subgraph of the AOP network containing the events enriched by the 31 engineered nanomaterials (ENMs) in the three biological systems: mouse lungs (orange), BEAS-2B cells (blue), and THP-1 cells (teal). Event type has been reported in the legend.

Mapping patterns of molecular alterations to the AOP has been shown to highlight similarities across biological systems [14]. This is also observed in the present study, where a core portion of the AOP network is enriched by all the test systems, covering events related to oxidative stress, NADH-ubiquinone and ROS production, TGF-β and cytokine release. The OECD has recently concluded that nanotoxicity related to human health mainly arises from inflammation, oxidative stress, and cytotoxicity (https://one.oecd.org/document/ENV/CBC/MONO(2022)3/en/pdf). These patterns were overlapping across test systems, suggesting that some early KEs involved in the initial assessment of hazard for ENMs, can indeed be tested *in vitro* while being representative of the *in vivo* counterpart. However, our results also show that test systems enrich different portions of the AOP network, *de facto* limiting the range of the assessable response mechanisms. THP-1 exposures specifically enriched pathways related to immunity and adipogenesis. Furthermore, the airway hyperresponsiveness is a specific mechanistic chain visible only in this cell line, as well as infertility and placental insufficiency. BEAS-2B enriched KEs associated with neuronal degeneration, lysosomal dysfunction, as well as events specific to epithelial cells. Importantly, BEAS-2B enriched lung fibrosis and collagen deposition, which is a relevant long-term consequence of ENM exposure, but is not visible in all the test systems. Overall, molecular events such as nuclear receptor activations appear to be better observable *in vitro*. Surprisingly, the chain of NFkB related KEs, a pivotal element of ENM-induced responses, are only enriched by *in vitro* systems. Exposures in the lung, on the contrary, enriched calcium homeostasis related pathways, and the complete mechanistic chain connecting nuclear receptors with PPAR gamma and adipogenesis. Internalisation of the compounds and DNA damage associated KEs are extensively observable in the lung, and partially shared with BEAS-2B. Given the experimental design of this dataset [27,28], this is a strong indicator that the differences in the observed events emerged because of the selected test system. The test system impact on the emerging phenotype upon compound exposure highlights the importance of considering both the exposed system and the compound in toxicological settings. A similar concept has been addressed by regulatory agencies as “the appropriateness of test species” [64]. However, this often only refers to the species-specific effects of ENMs. In contrast, we show here how the molecular buildup of different test systems impacts both the observed toxic effect and the detectable MOA in terms of toxicity testing.

Importantly, our results show that while some core events of ENMs exposure response are observable both *in vitro* and *in vivo*, the choice of the test system impacts the overall assessable response. While the applicability domain of a test system typically refers to the types of chemicals that can be tested, the present results suggest that exposures are not defined exclusively by the intrinsic features of the test substance. Our study indicates that i*n vitro* systems can better assess molecular events, but they may fail to capture entire mechanistic chains, which are observable in the *in vivo* setting. These considerations suggest that our framework could be used to contextualise existing molecular toxicological evidence with respect to the biological system. The impact of the molecular buildup of the test system informs on the consistency and biological concordance criteria which are usually evaluated to weight the relevance of omics data for regulatory hazard assessment.

## Conclusions

The ultimate aim of the 21^st^ century chemical assessment is to build a mechanistic understanding of the chemical-biological interactions. This can be achieved by using the AOP framework, including knowledge concerning MIEs and KEs with their underlying data, to produce models able to associate unstudied substances with known AOPs mechanisms [65]. In this study, we presented a novel network-based framework for linking toxicogenomics data to AOPs and for interpreting the response mechanisms of biological systems to chemical exposures. This framework acknowledges the inherent variability in how different biological systems respond to the same stimuli (exemplified here by ENMs) due to their unique molecular buildup. It enables the conversion of molecular alterations into chains of events across different levels of organismal complexity, ranging from molecular to tissue/organism events. As such, this approach provides a comprehensive interpretation of toxicogenomics data by exploiting the connections between toxicologically relevant events and gene interactions. To validate our framework, we conducted an extensive analysis of 93 exposures to 31 ENMs in three distinct biological systems. This dataset was selected for its comprehensive coverage of industrially relevant ENMs with varying physicochemical characteristics and well-defined hazard levels [11]. We systematically compared the outcomes of our network-based approach with traditional MOA investigations. Our results revealed that the network-based approach provided a deeper insight into the observed response. Representing MOAs as networks offers a framework to interpret multiscale complex events, extrapolating substantial more information as compared to other analytical frameworks for interpretation of toxicogenomics data. While network-based approaches are already well established in other fields of life sciences, systems toxicology applications are still in their infancy, with network properties hardly being exploited to solve toxicologically relevant issues [9,10]. This study underscores for the first time the potential of network theory in improving toxicogenomics data interpretability and its contribution to hazard assessment.

We also addressed the crucial issue of hazardous materials identification, a challenging task that traditional toxicogenomics methods struggle with. By applying our framework to exposures inducing various levels of toxicity, we highlighted common features and distinct molecular response mechanisms in each group. This approach allowed us to connect highly hazardous materials with molecular alterations associated with more apical events in the AOP framework. Importantly, this suggests that toxicity could be re-defined as the ability of inducing a multiscale alteration reaching a new stable patho/physiological state (e.g., an AO) in a particular experimental setting. In this light, no exposure would be “inert”; instead, it simply induces an alteration that can be easily reverted to homeostasis. This key insight, and the link to the molecular layers, are important inputs to the future design of quantitative AOPs.

Importantly, we showed that the choice of the test system profoundly influences the observed response mechanism, emphasising the need to consider the interaction between the compound and the biological system when assessing hazards. This study also highlights the importance of the molecular buildup (defined by the genotype and epigenotype) of different test systems in toxicological settings and its impact on the observable toxic effects and mechanisms of action. From a practical study design point-of-view, this means that toxicologists need not only to describe the features of the test materials and model systems [66] but also need to consider whether biological responses are assessable or not given the choice of the model system [2].

In conclusion, our network-based framework represents a significant step forward in understanding the complex relationship between chemical exposures and biological responses. It has the potential to contribute to toxicological assessments by providing a more comprehensive and informative perspective on toxicogenomics data interpretation. As the biomedical knowledge encoded in the AOP framework will increase, this approach will be also potentially useful to further characterise pathophysiological mechanisms of disease. Our observations could be seamlessly integrated into future safe-and-sustainable-by-design frameworks, offering a powerful tool for researchers and regulators alike. This framework overcomes some limitations of current omics based approaches, increasing interpretability and extrapolation of the effect of the specific exposure components, while also representing the first example of the potential contribution of systems and network theory to the hazard assessment of ENMs and, potentially, of other advanced materials.

## Methods

### Data

#### Datasets selection and preprocessing

The case studies are based on the publicly available datasets GSE148705 and GSE157266. These datasets we previously generated cover a broad panel of ENMs including metal, metal oxide, and carbon-based ENMs with different core (Ag, Au, TiO_2_, CuO, nanodiamond and multiwalled carbon nanotubes) and surface chemistries (COOH, NH_2_, or PEG). Each material was tested at equipotent doses (EC_10_) in the two different human cell lines (THP-1 and BEAS-2B), and at a dose of 10 µg/day for four consecutive days in mice. For further details, refer to Gallud et al. [27] and Kinaret et al. [28].

Transcriptomic data were preprocessed with the EUTOPIA software as previously described [6,67]. For the GSE157266 dataset, mouse genes have been converted to their orthologs, excluding the genes with no 1:1 relationship between mouse and human genome. In these datasets, for each ENM exposure, three replicates were present.

The final number of genes shared among all the datasets were 11972. In the preprocessing phase, the MDS highlighted outlier samples both in the BEAS-2B and Mouse datasets. Therefore the samples ‘RNA_2_NTND_24_0_1’, ‘RNA_2_NTQD_24_0_1’, ‘RNA_2_NTCuO_24_0_1’, ‘RNA_2_NTND_24_0_3’ and ‘NS6_1_4’ have been discarded from the datasets, respectively.

#### AOP network and annotation

In order to link the transcriptomic alteration induced by ENMs exposure, we exploited the AOP-gene annotation curated in [14]. The gene sets associated with each KE in the AOP were previously described in [23] The AOP network used in the framework was obtained by exploiting the connections between KEs present in the AOP-Wiki database (https://aopwiki.org/, downloaded on 26.10.2022).

### Computational framework

#### Framework input: Network inference and differential analysis

In the current study, input data for the framework were either in the form of a list of genes to be enriched or of a network (Figure 2).

Differential expression analysis was performed using the “limma” R package to compute the gene expression difference between exposed and controls, and correcting the P-value using the Benjamini– Hochberg. The normalised and corrected expression matrix was filtered according to standard thresholds (p-value < 0.05 and |logFC| > 0.58).

In order to infer co-expression networks for each exposure, we first defined gene nodes that would be shared in all the transcriptomic profiles. We selected genes present in all the studied dataset, where for human and mouse a conversion between orthologs has been performed, and mapped them to Ensembl ID. For each individual exposure, we ranked the genes based on both fold change and p-values, and selected the 100 top and bottom deregulated genes. The union resulted in 3061 genes.

For each pair of genes, we performed a Hotelling test (using the R package Hotelling, version 1.0), to identify differences in their multivariate means [68]. Edges were ranked according to their p-value from the lowest to the highest. Only edges with a p-value lower than the 5% percentile of the distribution were included in the network. This was repeated for each triple resulting in 93 binary networks. Information on the p-value was stored as an edge attribute in the exposure specific network (log), while information on the fold change was stored as node attribute and used in further steps of the analysis for filtering.

#### AOP gene sets enrichment and topological enrichment

In order to obtain enriched KEs for input lists of genes, we performed an enrichment of the differentially expressed genes. Differential expression analysis was performed as described under the *“Framework input: Network inference and differential analysis”* section. Then, enrichment was performed against KEs associated gene sets using Fisher’s exact test implemented in the *enrich* function of the R package “bc3net” [69]. Gene sets of each test system and genes from clustered KEs were merged, and unique genes were used to prepare the background of the enrichment.

In case a network is used as input for the framework (both in the cases in which we analysed a co-expression network or overrepresented set of edges) a topological enrichment was developed and used to find overrepresented KEs in the AOP network. First, all shortest paths between all nodes on the input network were estimated with the sknetwork.path.get_shortest_path function [70]. For each gene set (comprising the genes associated with a given KE) the average shortest path between each gene pair on the co-expression network was calculated. 1000 t-tests were performed by comparing the average shortest path of the known gene set versus randomly drawn gene sets of the same size. After estimating the average p-value, a key event was considered to be enriched if the average shortest path length of the known gene set divided by the average shortest path length of the random gene set was lower than 1.

#### Reconstruction of the AOP based mechanism of response

In order to represent the MOA of the exposure as a network of KEs, we created a framework which filters the most relevant KEs, prioritising “plausibile” MIEs and AOs, and finally maps it back to the AOP network.In this context, plausibility was defined both as a measure of proximity to an enriched KE on the AOP network, as well as the presence of statistically altered genes in the annotated gene set of the event.

First, the enrichment p-values were corrected using the “fdr” method with the p.adjust function of the R package “stats”. KEs with more than 1000 genes annotated and adjusted p-value higher than 0.05 were discarded. On the filtered results, only events with a p-value lower than the 5-th percentile of the distribution were selected. Finally, KE where genes annotated had a fold change lower than 0.58 were discarded.

Second, probable MIEs and AOs were prioritised based on the expression of the annotated genes, the closeness on the AOP network, and the size of the annotated gene set. Closeness was computed with the shortest.paths function in the “igraph” R package [71]. Events with no annotated gene differentially expressed were discarded. MIEs and AOs were ranked based on the p-value of the expression of the annotated genes and the distance from the enriched KE. In case more than 20 events were prioritised, 20 was considered as an arbitrary cut-off. All the prioritised events were mapped on the AOP network and plotted using the “igraph” R package [71].

### Network comparison

#### Definition of information measures

In order to quantitatively compare our framework results obtained using either co-expression networks or lists of differentially gene sets as an input, we defined two measures of information. First, we defined information as the capability of reconstructing an AOP based mechanism from a MIE to an AO, which we defined as the “AOP completeness score”. In this light, we computed, for each reconstructed network of ENM response, the transitivity, betweenness, shortest paths and diameter using the homonym functions of the igraph R package [71]. When multiple values were returned (e.g., betweenness and shortest paths), the mean of the values distribution was considered. These measures indicate a more connected network, thus a better possibility of reconstructing a unified mechanism of response between causally linked events. For this reason, we considered a greater (mean) value as an indication that a longer mechanism could be completed. When comparing the results of two mechanisms, we evaluated as more informative the response network having at least three out of four higher values of connectedness.

Second, we considered the possibility of having a more predictable or homogeneous mechanism, which can be a desirable gain as the information content is less uncertain. We defined it as The “Key event connectivity score”, namely the ratio between the portion of connected KEs against the unconnected ones (isolated nodes) characterising the MOA of an exposure.

#### Computation of a biological system similarity score between network responses

We investigated whether the obtained mechanism of response could be used to derive the similarities between exposures of the same material across biological systems. We defined this measure as the consensus of four difference parameters. First, we evaluated how similarly the two networks cover a portion of the complete AOP network. For this, we considered the delta of the ratio between each network number of KEs to the complete AOP.

Second, we considered the overlap between the two response networks, computing a jaccard index of the KEs nodes and edges.

Finally, we computed the ratio of the diameters of the biggest component (computed with the diameter function in the R package “igraph”), as it represents the largest retrievable mechanism in each response network [71]. The mean of all these scores was used as a similarity measure between the networks.

### Network analysis

#### Clustering strategies for co-expression networks and AOP events networks

Co-expression networks were clustered using the functions developed in VOLTA, a python package for co-expression network analysis [72]. Briefly, we first computed edge based similarities between each network pair using the get_edge_similarity function. Then the consensus_clustering function was used based on the three individual clusterings (hierarchical, affinity propagation, k medoids).

In order to cluster the AOP based mechanisms of response, we first computed a similarity score as the mean of the jaccard index between networks nodes and edges. This similarity reflects the number of shared nodes and edges between the response mechanisms. For each pair of networks, the similarity was converted in distance. Finally, the clustering was performed using the “hclust” function from the R package stats, with the ward.D2 method, and results plotted as a dendogram.

#### Identification of overrepresented structures

In order to highlight edges overrepresented in groups of networks, we exploited the function get_statistical_overrepresented_edges from VOLTA in the pattern matching module. Briefly, the function finds edges that are statistically overrepresented in a cluster (group of networks) based on a hypergeometric function and Benjamin-Hochberg correction. Edges associated with specific biological systems, nanoparticle surface modifications, and toxicity endpoints were retrieved in the same way.

#### Edge based enrichment analysis

Classical enrichment analysis is used to identify classes of genes that are over-represented in large sets. However, when performing enrichment analysis over the nodes co-expression network, the classical algorithm does not consider if the genes connected by edges in the network are part of the same pathways. In order to consider gene-gene association we performed enrichment of network edges against gene pairs derived from human pathways.

To create the background, we collected data for pathway gene sets in the KEGG databases [73]. Background edges were created based on the annotation of genes to the same gene set. Input edges were ranked from the most to the least co-deregulated, based on their p-value. In case of edges present in multiple exposures (like overrepresented edges), the individual p-values were combined using the sumlog method. The enrichment analysis was performed by means of a weighted Kolmogorov-Smirnov test, to identify edge sets that are over-represented on top of the rank. The statistical significance of the enrichment analysis was estimated by permutation analysis over 100 random shuffles of the edge sets. The p-values were corrected for multiple comparisons using the false discovery rate (FDR) method and setting 0.05 as the significance threshold. The analysis was performed by using the code originally developed in [74].

## Data and code availability

The transcriptomic dataset used for the case studies of this analysis are freely available at GEO (https://www.ncbi.nlm.nih.gov/) under accession numbers GSE146708, GSE148705, GSE157266. Preprocessed data has been previously deposited also in the online Zenodo repository (https://doi.org/10.5281/zenodo.7674574). The AOP annotation is freely available in the online Zenodo repository (https://doi.org/10.5281/zenodo.7980953)[23]. KEs and KE relationships information have been downloaded from https://aopwiki.org/ on 26.10.2022. KEGG functional data were downloaded on 12.2019 and curated as described in [75]. All the other relevant data and data supporting the findings of this study have been deposited in the online Zenodo repository available at https://doi.org/10.5281/zenodo.10390383. All the relevant and custom code supporting the findings of this study has been deposited in the online Zenodo repository https://doi.org/10.5281/zenodo.10390383.

## Author contribution

**Giusy del Giudice:** Data curation, Formal analysis, Investigation, Validation, Methodology, Software, Visualization,Writing - original draft, Writing - review & editing. **Angela Serra:** Formal analysis,Validation, Methodology, Software, Supervision,Writing - original draft, Writing - review & editing. **Alisa Pavel:** Methodology, Software, Validation, Writing - review & editing. **Marcella Torres Maia:** Formal analysis,Writing - original draft, Writing - review & editing. **Laura Aliisa Saarimäki:** Investigation,Writing - original draft, Writing - review & editing. **Michele Fratello:** Methodology, Software, Writing - review & editing. **Antonio Federico:** Supervision, Writing - review & editing. **Harri Alenius:** Investigation,Writing - review & editing. **Bengt Fadeel:** Investigation,Writing - review & editing. **Dario Greco:** Conceptualization, Supervision, Funding acquisition, Project administration, Resources, Writing - original draft, Writing - review & editing.

## Acknowledgements

This study was supported by the Academy of Finland project UNICAST NANO (322761), by the European Research Council (ERC) programme, Consolidator project ARCHIMEDES (101043848), and the European Community’s Seventh Framework Programme (FP7) under Grant Agreement No. 309329 (NANOSOLUTIONS). A.S. and A.F. were supported by the Tampere Institute for Advanced Study.

